# Intraoperative enrichment of bone autograft with autologous bone marrow-derived mononuclear cells isolated simultaneously: moving towards tissue engineering *in situ*

**DOI:** 10.1101/2020.11.09.375188

**Authors:** D.S. Baranovskii, B.G. Akhmedov, O.A. Krasilnikova, A.G. Demchenko, M.E. Krasheninnikov, M.V. Balyasin, O.Yu. Pavlova, N.S. Serova, I.D. Klabukov

## Abstract

**Background:** The use of tissue-engineered bone autografts is a promising approach for bone defects restoration. The isolation of cells and their seeding on bone autograft is usually carried out in a laboratory, requiring significant time and two separate surgical interventions. Intraoperative creation of tissue-engineered bone autograft can represent a perspective solution. The aim of this study is to investigate the possibility of creation of tissue-engineered bone autograft by intraoperative enrichment of bone tissue with bone marrow-derived mononuclear cells (BM-MNCs) isolated simultaneously.

**Methods:** Red bone marrow and autologous bone tissue (bone fragments and bone chips) of the donor were harvested intraoperatively. BM-MNCs were isolated, and bone fragments were enriched with BM-MNCs intraoperatively. Assessment of the adhesion and proliferation of BM-MNCs on bone fragments was carried out by fluorescence microscopy and histological examination. MTT assay was used to compare metabolic activity of BM-MNCs and wBMA cells seeded on bone chips.

**Results:** Autologous bone fragments were colonized with autologous BM-MNCs isolated simultaneously in the O.R. with further adhesion and active growth of cells. When seeded on bone chips, metabolic activity of BM-MNCs was statistically significantly higher compared to wBMA cells (p-value=0.0272) on day 14. There was no difference in metabolic activity of BM-MNCs and wBMA cells cultured in nutrient medium without bone chips.

**Conclusion:** Technically simple method of intraoperative enrichment of autologous bone fragments with BM-MNCs isolated simultaneously allowed to create tissue-engineered bone autograft in the O.R. The safety and effectiveness of intraoperatively enriched autografts should be investigated further.

## Introduction

A successful repair of extended bone defects is one of the leading challenges of orthopedics and traumatology. In particularly complicated cases localization of the defect does not allow the application of distraction osteogenesis, meanwhile spontaneous recovery of lost bone tissue is impossible due to the significant size of the defect. Currently, bone-graft autotransplantation is a “gold-standard” approach for such clinical cases. However, a well-known method of autologous osteoplasty is commonly limited by the bone volume that can be safely harvested from the donor area. Furthermore, insufficient vascularization and inflammatory reaction are often observed in the zone of bone regeneration^[1]^.

Previously published studies have shown that bone marrow-derived mononuclear cells (BM-MNCs) transplantation can stimulate the regeneration of bone tissue and increase its density and vascularization^[2–5]^. The regenerative potential of BM-MNCs is explained by the presence of different subpopulations of cells in the mononuclear fraction of bone marrow. For example, bone marrow-derived mesenchymal multipotent stromal cells, which are part of the mononuclear fraction of bone marrow, possess the ability for local suppression of inflammation, differentiation into osteoblasts, expression of paracrine factors, and stimulation of angiogenesis in the implantation zone^[6–7]^. In addition, clinical studies have shown that CD34+ cells, which also are BM-MNCs, exert a positive effect on bone tissue regeneration, increasing the activity of cells involved in bone regeneration through paracrine and autocrine interaction^[8]^. Therefore, enrichment of bone autograft with autologous BM-MNCs is capable of enhancing the regenerative potential of a bone transplant. BM-MNCs can be easily isolated from the bone marrow aspirate using gradient centrifugation technology. The automated cell separation system Sepax S100 allows obtaining a cell suspension intraoperatively with BM-MNCs concentration 3-10 times higher than that in the bone marrow aspirate^[9]^. The separation is carried out in a closed sterile single-use device excluding the potential risk of contamination.

Notably, bone marrow aspirate is also used for the enrichment of bone tissue and the enhancement of its regenerative potential^[10–11]^. The therapeutic potential of bone marrow aspirate is mainly attributed to the presence of endothelial progenitor cells and MSCs. In contrast to the BM-MNCs, bone marrow aspirate contains not only BM-MNCs, but also red blood cells, platelets, and their precursors, as well as leukocytes. However, in order to investigate the properties of adhesive cell populations *in vitro*, red blood cells and platelets within the harvested bone marrow are lysed, resulting in the washed bone marrow aspirate (wBMA)^[12]^.

Usually, isolation and seeding of BM-MNCs on the patient’s autologous bone material are carried out in the laboratory under aseptic conditions corresponding to GMP standards and require a considerable time and economic expenses. In addition, the patient undergoes two or more separate surgical interventions necessary for the extraction and implantation of autologous material. Moreover, in a number of countries, the administration of cultured cells is strictly regulated by biomedical laws that require registration of a cell product before clinical use^[13–15]^. Isolation of autologous BM-MNCs and seeding of the bone autograft with cells directly in the O.R. with simultaneous implantation of enriched autologous material to the patient may not only reduce the number of surgical procedures, time and cost of cell preparation, but also falls under less strict legal regulation.

The aim of the present study is to investigate the possibility of intraoperative creation of tissue-engineered bone autograft by the colonization of autologous bone material with simultaneously isolated BM-MNCs and to compare the efficacy of autologous bone chips colonization with BM-MNCs and wBMA cells based on the measurement of metabolic activity.

## Materials and methods

### Harvesting of red bone marrow and autologous bone material (bone fragments and bone chips)

Red bone marrow and autologous bone material for this study were harvested intraoperatively during bone autoplasty of post-traumatic ram bone defect in a 48-years-old male donor, who gave informed consent.

To extract bone autograft from the donor area, a skin incision was made 1 cm above the iliac crest. Periosteum, external, and internal cortical plates were dissected and filed with the surgical saw. The bone autograft was finally harvested using a chisel and a surgical clamp. The red bone marrow and bone material were collected through the surgical wound at the donor site of the iliac crest. In order to prevent coagulation, heparin was added to the bone marrow aspirate at the rate of 100 IU per 1 ml of aspirate. Harvested bone autograft was customized to achieve an optimal shape for implantation in the area of post-traumatic ram bone defect.

Four large autologous bone fragments with sizes from 0.5 to 0.8 cm were separated from the harvested bone autograft and were used in further studies in vitro. In addition, small bone pieces with diameters of 0.2 to 0.3 cm were crushed to produce bone chips, which were further seeded with BM-MNCs or wBMA cells for assessment of cell metabolic activity. The removal of the donor’s bone fragments and bone marrow for the study did not lead to an extension of the surgical procedure and did not require changes in the operation protocol.

### Isolation of BM-MNCs from the bone marrow aspirate

The donor’s BM-MNCs isolation was performed under aseptic conditions in the O.R. by gradient centrifugation in a sterile closed environment using the automated cell separation system Sepax S100 (Biosafe, Switzerland) in accordance with the standard protocol. The quantification of living cells in the obtained mononuclear fraction was carried out using the TC10 automated cell counter (BioRad, USA) by counting the number of living cells in the 20 μl of the suspension.

### Enrichment of autologous bone fragments with BM-MNCs in the O.R

Enrichment of the autologous bone fragments with BM-MNCs was carried out under aseptic conditions in the O.R. Briefly, 1 ml of the BM-MNCs suspension was added to each of the four bone fragments with subsequent incubation for 20 min at 37°C. The enriched bone fragments were placed in saline and transported to the laboratory in a hermetic container for further investigation together with the donor’s wBMA and suspension of BM-MNCs (Fig. 1).

**Figure 1.**
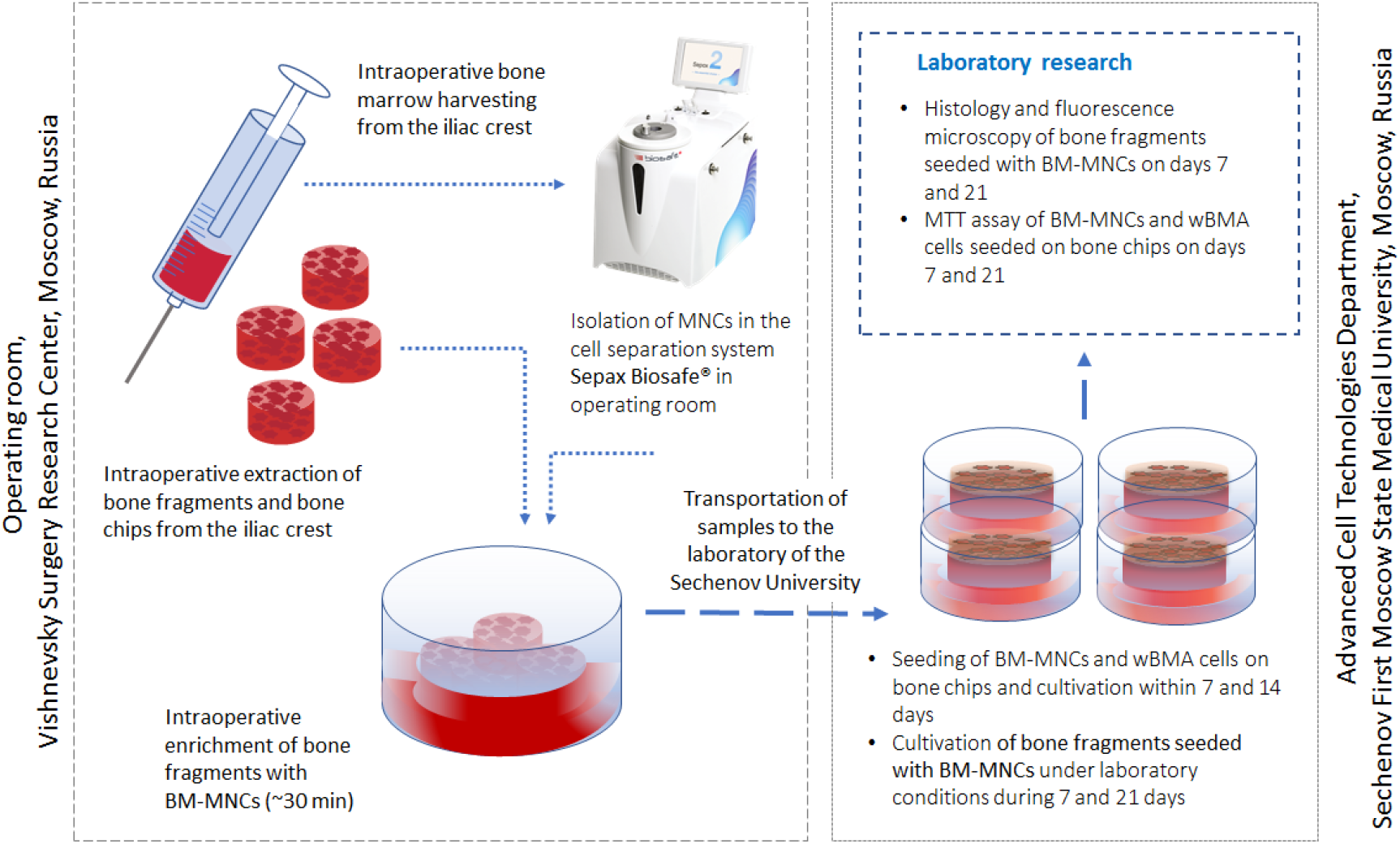
Intraoperative enrichment of autologous bone fragments with simultaneously isolated BM-MNCs and comparison of metabolic activity of BM-MNCs and wBMA cells, seeded on bone chips.

### Cultivation of BM-MNCs on autologous bone fragments

Cultivation of BM-MNCs on autologous bone fragments was carried out under laboratory conditions after the transportation of enriched autologous bone fragments from the OR. Cultivation of BM-MNCs on enriched bone fragments (n = 4) was carried out for 7 and 21 days in a 12-well plate in the nutrient medium containing DMEM medium (Gibco, USA), 10% of fetal bovine serum (Gibco, USA), 100 units/ml of penicillin and 100 μg/ml of streptomycin (Gibco, USA) in a 5% CO_2_ incubator at 37°C.

### Histological examination

For histological examination on day 7 after cell seeding, a 2×2 mm sample was taken from the surface of each bone fragment. For histological examination at day 21 after cell seeding, enriched bone fragments remaining after sampling were used. Briefly, samples and bone fragments were fixed in 10% formalin, dehydrated in a graded series of alcohols, and embedded in paraffin. Sections were stained with hematoxylin and eosin (H&E) according to the standard protocol for bone tissue and analyzed using a light microscope Nikon L200 (Nikon, Japan) equipped with a fluorescent illuminator and a digital camera.

### Fluorescence microscopy

To study the adhesion and proliferation of BM-MNCs on autologous bone fragments, fluorescence microscopy of the surface of all four autologous bone fragments was performed on days 7 and 21 after cell seeding. Briefly, one 2×2 mm sample was taken from the surface of each bone fragment. Samples were examined using a light microscope Nikon L200 (Nikon, Japan) with RNase A pre-treatment and staining of cell nuclei with ethidium bromide using a standard protocol.

### Comparative study of metabolic activity of BM-MNCs and wBMA cells seeded on bone chips in vitro

Evaluation of the metabolic activity of BM-MNCs and wBMA cells seeded on bone chips or cultured in the nutrient medium was performed by MTT assay. The bone chips obtained intraoperatively from bone fragments were evenly distributed in 20 wells of two 96-well plates. Untreated heparinized wBMA cells and the BM-MNCs were centrifuged separately for 5 min at 1500 rpm. The cell pellet was diluted in a lysis buffer containing 114 mM NH_4_Cl, 7.5 mM KHCO_3_, 0.1 mM EDTA. Next, the suspension was centrifuged and the cell pellet was diluted in a complete nutrient medium. The number of living wBMA cells and BM-MNCs in the suspension was detected using automatic cell counter Countess (Invitrogen, USA). Next, 10 μl of the suspension of wBMA cells and BM-MNCs per well were placed in a 96-well plate containing bone chips (10 wells for each type of suspension). For adhesion, the cells were left in an incubator (37°C, 5% CO_2_) for 30 minutes, then 100 μl of the nutrient medium was added to each well. As a positive control, 10 wells with BM-MNCs and 5 wells with wBMA cells were used; a nutrient medium was used as a negative control. Every two days red blood cells were removed with Hank’s Balanced Salt Solution and the nutrient medium was changed. Cell viability was assessed on days 7 and 14 using MTT assay. Briefly, the nutrient medium was changed and 250 μl of 0.48 mM MTT reagent were added to each well and incubated (37°C, 5% CO_2_) for 4 hours. Next, the supernatant was removed and the formed formazan crystals were dissolved in 150 μl of DMSO. Absorption was measured at a wavelength of 540 nm on a Multiskan FC (Thermo) photometer. The background value was taken into account by adding the reagent to the nutrient medium (n=3).

### Statistics

Statistical analysis was performed in the GraphPad Prism 7.0 software (USA). One-way ANOVA was used for multiple comparisons. The threshold of statistical significance was established as a p-value<0.05.

### Ethical statement

Written informed consent was provided by the participant of this study.

## Results

### Viability of BM-MNCs in the mononuclear fraction of bone marrow aspirate

The automatic separation allowed us to obtain 19 ml of a ready-to-use suspension of BM-MNCs from 80 ml of harvested bone marrow aspirate in 30 minutes. The automatic counting of cells in the isolated mononuclear fraction showed a high concentration of viable BM-MNCs in the suspension - not less than 260×10^3^ cells/ml.

### Adhesion and proliferation of autologous BM-MNCs on autologous bone fragments

Fluorescence microscopy of enriched autologous bone fragments showed a substantial number of adhered and fluorescenting BM-MNCs on the surface of bone material on days 7 and 21 after cell seeding.

On day 21 (Fig. 2b) the number of BM-MNCs on the surface of the bone fragments significantly increased compared to day 7 (Fig. 2a) suggesting active proliferation of BM-MNCs.

**Figure 2.**
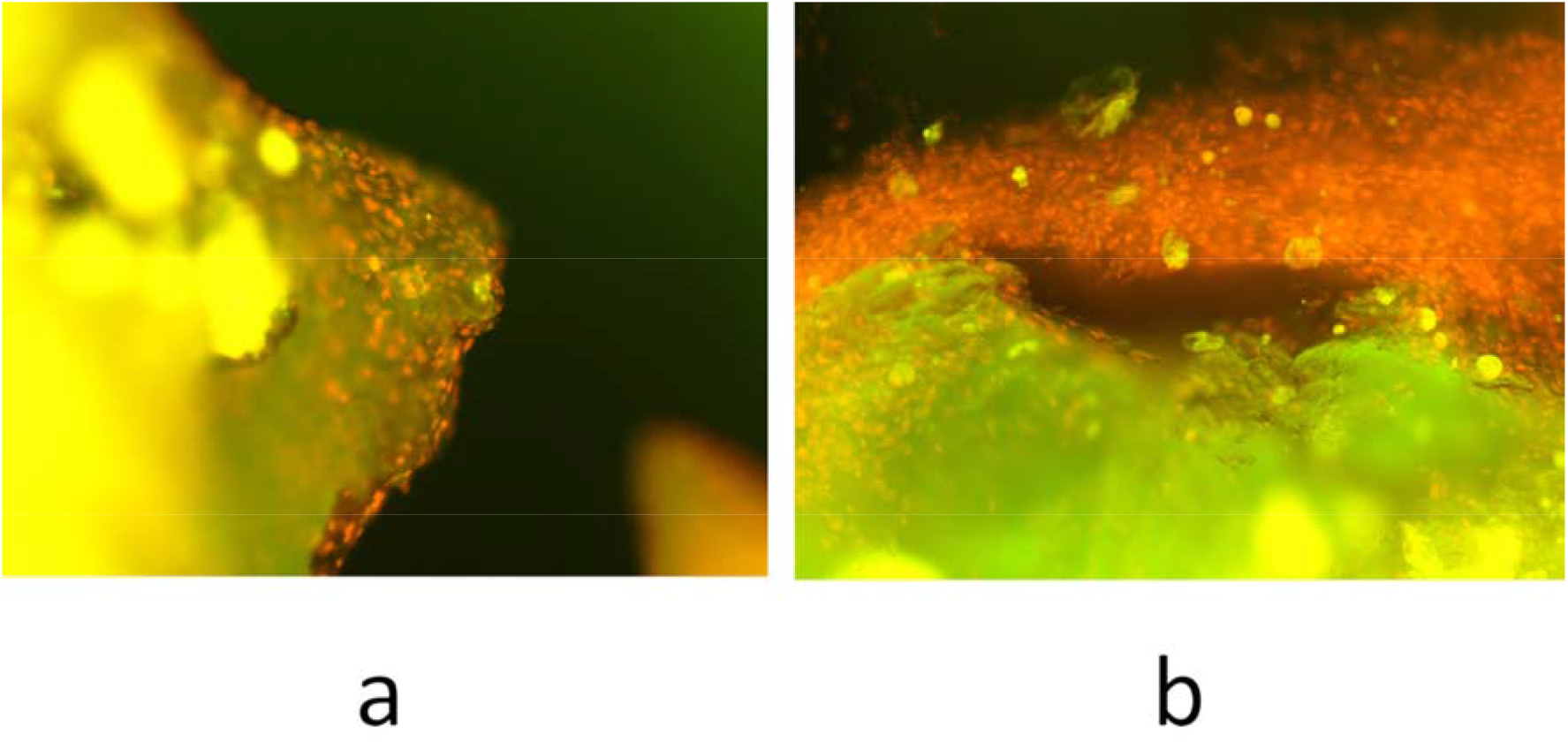
Autologous bone fragments enriched with autologous BM-MNCs: (a) 7 days after seeding, (b) 21 days after seeding. Fluorescence microscopy, orange fluorescence of BM-MNCs nuclei. Ethidium bromide staining with RNase A, x200 magnification

### Histological study of autologous bone fragments enriched with BM-MNCs

Histological study showed the presence of viable BM-MNCs on the surface of autologous bone fragments on days 7 and 21 after cell seeding (Fig. 3).

**Figure 3.**
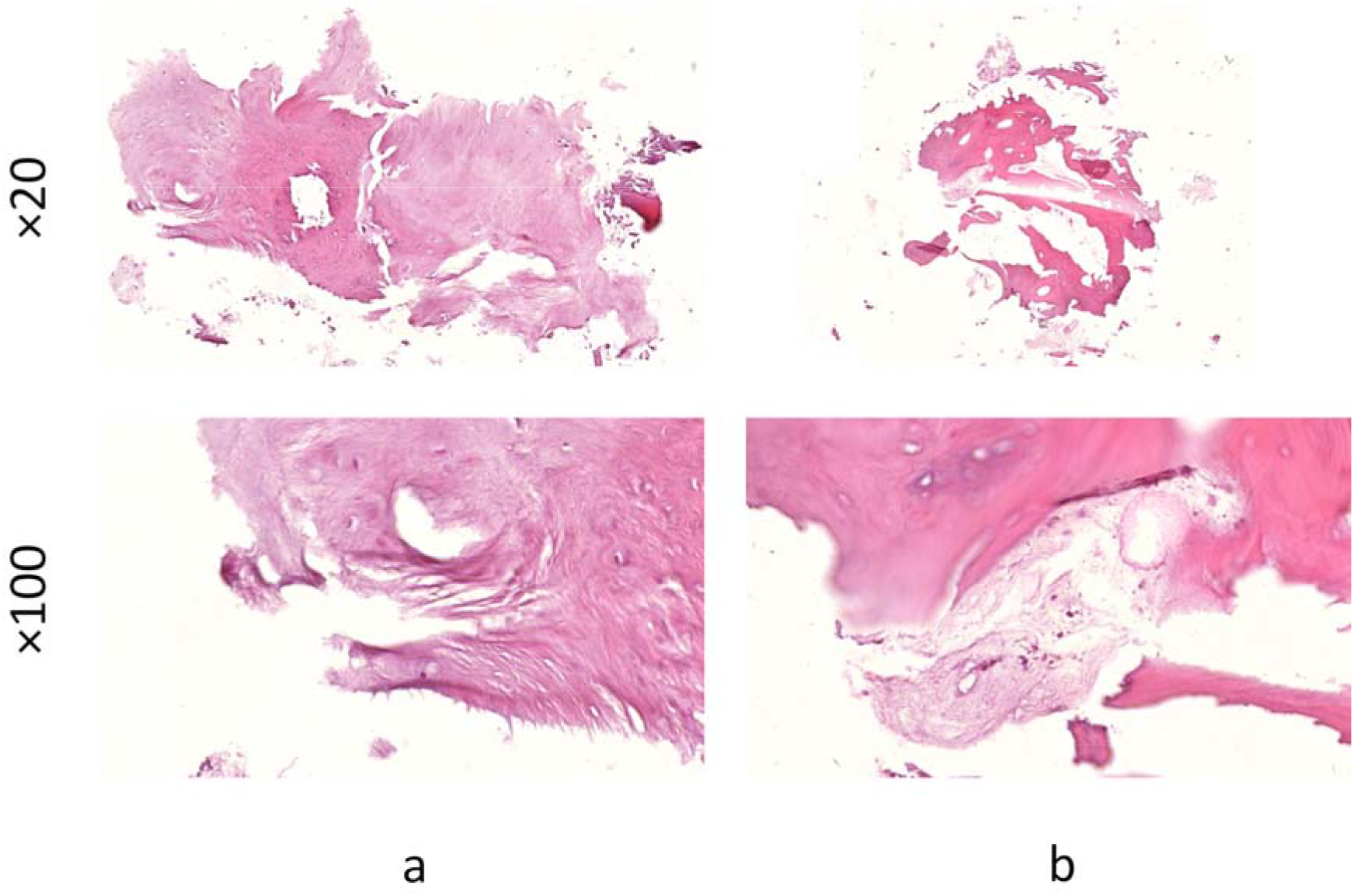
Autologous bone fragments enriched with BM-MNCs, H&E staining, light microscopy: (a) day 7 after seeding, (b) day 21 after seeding

Cell clusters and separate BM-MNCs with dark nuclei stained with hematoxylin are observed on the surface of eosinophilic bone tissue, covering its surface with a thin layer on day 7 after seeding (Fig. 3a). The newly formed BM-MNCs layer grows over time and is clearly visualized during histological examination on day 21 after seeding (Fig. 3b).

### Metabolic activity of BM-MNCs and wBMA cells seeded on bone chips

The number of viable wBMA cells and BM-MNCs seeded on bone chips was about 200 × 10^3^ per well. The results of the evaluation of the metabolic activity of cells seeded on bone chips on days 7 and 14 performed by MTT assay are shown in Figure 4.

**Figure 4.**
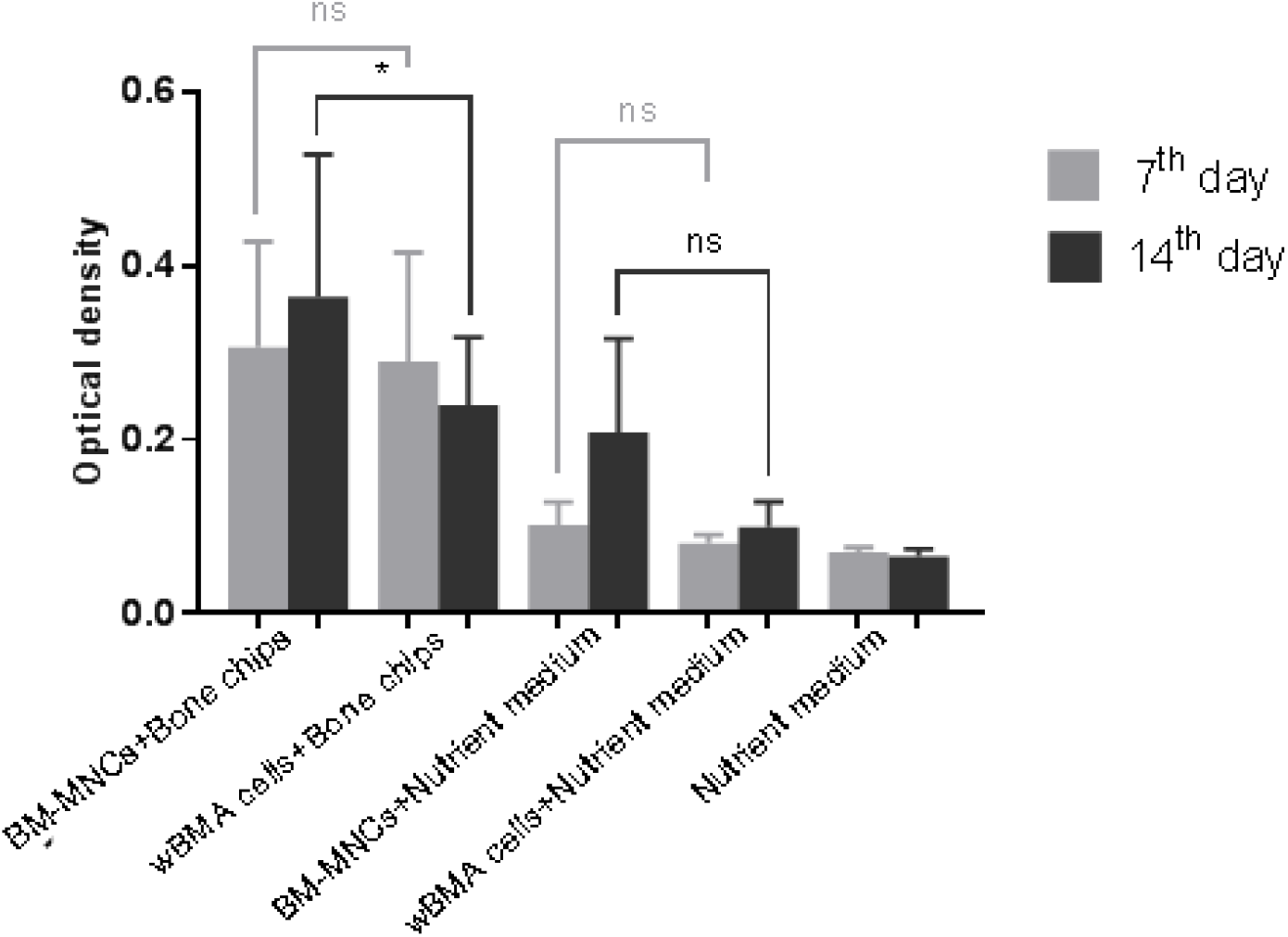
Results of the evaluation of cells’ metabolic activity on days 7 and 14 after enrichment of bone chips with BM-MNCs or wBMA. Negative Control: cell-free nutrient medium. *: p-value<0.05, ns: not significant

Statistically significant differences in the optical density of MTT-formazan on day 14 were revealed between the groups of BM-MNCs with bone chips compared to wBMA cells with bone chips (p-value=0.0272), while between groups of BM-MNCs+nutrient medium compared to wBMA+nutrient medium on days 7 and 14 the differences were not statistically significant (p-values>0.05).

## Discussion

Recent studies have shown that the use of tissue-engineered bone grafts in clinical practice can represent a promising approach for the effective restoration of bone defects^[16–18]^. However, the manufacturing of tissue-engineered bone autografts usually requires time-consuming cultivation of cells in vitro and further enrichment of the bone autograft. At the same time, intraoperatively created tissue-engineered bone autografts may help to reduce time, expenses, and a number of surgical interventions. In the present study, we have shown the possibility of intraoperative isolation of BM-MNCs fraction with simultaneous enrichment of autologous bone material directly in the O.R. Notably, fully automated separation allows to achieve a safe and standardized BM-MNCs isolation process. At the same time, the short period required for the separation of BM-MNCs allows performing the procedure without extending time of the surgery. Importantly, time for the adhesion of MB-MNCs on the bone fragments was 20 minutes, and neither caused extension of surgery time. Further studies, carried out under laboratory conditions, demonstrated colonization of autologous bone fragments with BM-MNCs, and active proliferation of cells on bone fragments during 14 days.

Notably, not only BM-MNCs but also bone marrow aspirate cells seeded on implants are capable of improving bone tissue regeneration in humans^[10,19,20]^.

In the present research, it has been shown that the metabolic activity of BM-MNCs seeded on bone chips was higher than that of wBMA cells. The difference in metabolic activity may be due to the fact that mononuclear fraction contains a higher percentage of dividing cells, while leukocytes in the wBMA do not proliferate. However, it is noteworthy that there was no difference in metabolic activity between BM-MNCs and wBMA cells cultivated in the nutrient medium without bone chips. Consequently, we speculate that adhesion to bone chips has a stimulating effect on the metabolic activity of BM-MNCs compared to wBMA cells and may be explained by specific adhesion of BM-MNCs to bone tissue.

Since the correlation of optical density in MTT assay with the number of cells has been shown previously^[21]^, it is possible to conclude that BM-MNCs had not only a greater metabolic activity in comparison with wBMA cells, but also a greater proliferation rate. The results of the present study are in accordance with previously published data on the advantages of using mononuclear fraction in osteoinduction in comparison with wBMA cells after cultivation^[22]^.

The technical possibility of fast and automated isolation of BM-MNCs and their seeding on autologous bone material in the operating room together with obtained data on the higher metabolic activity of BM-MNCs in comparison with wBMA cells allows us to suggest that the described method can be used for intraoperative creation of tissue-engineered bone autografts that can contribute to the more effective repair of bone damage.

## Conclusion

In the present study, a technically simple method of intraoperative enrichment of autologous bone fragments with BM-MNC isolated simultaneously was described. It was demonstrated that intraoperatively isolated BM-MNCs have the ability to attach and proliferate on the surface of the autologous bone material. The use of automated separation devices for isolation of autologous BM-MNCs allows creating tissue-engineered bone autograft directly in the O.R., reducing the number of surgical procedures for the patient, thus reducing the risks of postoperative complications. The safety and effectiveness of intraoperatively enriched bone autografts should be a subject of further preclinical and clinical studies.

## Acknowledgments

The authors would like to thank Mr. Lev K. Abdullaev, a laboratory assistant at the Advanced Cell Technologies Department of Sechenov University, for his help in organizing the experiment. This work was carried out using equipment of the Shared-Use Facility Center “Regenerative medicine” of Sechenov University (ID310020).

